# Predictable Modality Transitions and Amodal Representations Enable Crossmodal Statistical Learning

**DOI:** 10.1101/2023.05.12.540508

**Authors:** Alexis Pérez-Bellido, Daniel Duato, Francesco Giannelli, Ruth de Diego-Balaguer

## Abstract

Statistical learning involves extracting statistical regularities from the environment. Despite the ubiquitous crossmodal relations in our environment, previous research has failed in showing crossmodal learning suggesting that statistical learning is modality-specific, occurring within but not between sensory modalities. The present study investigates under which circumstances statistical learning can occur between modalities. In the first experiment, participants viewed a stream of meaningless visual fractals and synthetic auditory stimuli. Importantly, the sequence of stimuli could be grouped into unimodal or crossmodal pairs based on their internal transitional probabilities. Using implicit and explicit measures of learning, we found that participants only learned the unimodal pairs. In the second experiment, pairs were presented in separate unimodal and crossmodal blocks. The crossmodal blocks alternated visual and auditory modalities allowing participants to anticipate the upcoming modality. This manipulation allowed significant statistical learning for the crossmodal pairs, reflected only by implicit measures. This suggests that modality transitions predictability aids correct attention deployment across sensory modalities, crucial for learning crossmodal statistical contingencies. In the third experiment, where audiovisual stimuli with semantic content were used leaving access to an amodal shared representation, participants could explicitly recognize statistical regularities between crossmodal pairs even when the upcoming modality was unpredictable. This finding suggests statistical learning between crossmodal pairs can occur when sensory-level limitations are bypassed, and when learning unfolds at an amodal level of representation. These findings challenge the view that statistical learning is strictly modality-specific, instead indicating that crossmodal statistical learning depends on attentional mechanisms and representational level.

**Public significance statement:** Previous research has established that low-level statistical learning occurs within individual sensory modalities. However, evidence for statistical learning across modalities remains limited. This has led to the hypothesis that statistical learning operates as a modality-specific system, constrained by the perceptual properties and content of each sensory system. In this study, we demonstrate that modality transitions’ predictability enables crossmodal statistical learning, even in the absence of semantic information. Notably, this constraint does not apply to meaningful stimuli, which can be processed at an amodal level. Our findings broaden the classical neurobiological model of statistical learning by highlighting the roles of attentional mechanisms and representational levels in supporting crossmodal statistical learning.

## Introduction

Environmental information is highly structured conforming temporal and spatial patterns. The human brain has developed an exquisite sensitivity to implicitly detect these statistical regularities and exploit them to efficiently anticipate upcoming environmental changes. This ability, termed statistical learning, has been thoroughly investigated in the last decades showing that the human brain is ready to pick-up on statistical regularities at very early ages of development (Saffran et al., 1996). Statistical learning is evolutionarily rooted, as it has been observed in other animals, and it operates across sensory modalities, including auditory, visual, and tactile systems (Conway & Christiansen, 2005; Fiser & Aslin, 2001; Saffran et al., 1999). Concerning the relation between modalities, there is empirical evidence demonstrating that statistical learning in one sensory modality is not impervious to stimulation in other sensory modalities (Cunillera, Càmara, et al., 2010; Cunillera, Laine, et al., 2010; Glicksohn & Cohen, 2013; Mitchel & Weiss, 2011; Thiessen, 2010; Toro et al., 2005). However, the evidence showing that statistical learning takes place between sensory modalities is limited and inconsistent (Frost et al., 2015; Walk & Conway, 2016).

For instance, prior research has demonstrated that humans can concurrently learn statistical regularities independently in two sensory modalities without interference, pointing to parallel statistical learning mechanisms across perceptual systems (Conway & Christiansen, 2006; Li et al., 2018; Mitchel & Weiss, 2011; Seitz et al., 2007). Additionally, statistical learning of an artificial grammar does not transfer across modalities (Redington & Chater, 1996; Siegelman & Frost, 2015) and there is no empirical evidence showing that humans can detect statistical regularities between inputs presented in different sensory modalities (Walk & Conway, 2016). As a result, the most accepted framework in the literature postulates that while statistical learning relies on a domain-general set of computations, learning of the statistical associations between low-level sensory features is modular and constrained by the specific characteristics of each sensory modality (Frost et al., 2015). From this perspective, statistical learning of low-level sensory regularities would be modality-specific, taking place in their corresponding sensory areas through neural plasticity changes (Reber, 2013). Crossmodal statistical learning would be exclusively limited to those sensory inputs that encode high-level representations of meaningful stimuli, and in theory could occur through amodal representations shared between modalities.

In this non-preregistered study, we aim to scrutinize this framework in three experiments while testing three complementary hypotheses that might explain why finding statistical learning between low-level sensory inputs is so elusive. Firstly, studies on crossmodal statistical learning typically involve a phase in which participants are passively exposed to statistical regularities, followed by a phase in which participants must explicitly recognize the crossmodally associated items (Conway & Christiansen, 2005; Seitz et al., 2007; Walk & Conway, 2016). This approach has obvious limitations, as relying solely on explicit tests may provide an incomplete depiction of whether participants have learned or not. Statistical learning typically unfolds in parallel in both implicit and explicit fashions (Batterink et al., 2015; Conway, 2020), with the implicit system always “on” but the explicit system optional. Therefore, for completeness, implicit statistical learning must be quantified using an online measure of performance during the exposure phase (Siegelman et al., 2017). Indeed, previous studies utilizing implicit measures have contributed to uncovering statistical learning, as demonstrated by the facilitation in detecting expected targets or discriminating unexpected noisy stimuli during the learning phase (Barakat et al., 2013; López-Barroso et al., 2016; Orpella et al., 2021; Richter et al., 2018; Richter & de Lange, 2019). Thus, it is possible that previous studies that only assessed learning based on explicit statistical learning measures might have overlooked significant but implicit crossmodal statistical learning effects.

Second, crossmodal statistical learning might be impeded compared to unimodal statistical learning due to a modality switching cost. Specifically, in randomly intermixed sequences of multimodal trials, reaction times are slower when the target stimulus is preceded by a stimulus delivered in a different sensory modality (Gondan et al., 2004; Otto & Mamassian, 2012; Shaw et al., 2020). This phenomenon, defined as modality switching cost, is associated with an attentional bias where we tend to expect information in the last processed sensory modality (Spence et al., 2001). In previous studies investigating crossmodal statistical learning (Walk & Conway, 2016), multimodal stimuli were randomly intermixed during the learning phase. Therefore, the presence of modality switching costs could potentially introduce biases favoring the perceptual grouping of inputs delivered within the same sensory modality. As suggested in Conway (2020), while learning unimodal regularities may be relatively independent of attention, the acquisition of statistical regularities across different sensory modalities may rely more heavily on precise selective attentional mechanisms. Consequently, if such a bias in perceptual grouping exists, it could affect the ability to bind together crossmodal items and learn crossmodal transitions effectively.

Third, previous studies assessing crossmodal statistical learning (Seitz et al., 2007; Walk & Conway, 2016) used meaningless audiovisual stimuli. According to the framework proposed by Frost and colleagues (Frost et al., 2015), crossmodal statistical learning can only occur between amodal representations. Therefore, the hypothesis that the level of representation of unisensory inputs determines the formation of crossmodal associations remains to be tested.

To test these hypotheses, we designed three statistical learning experiments where participants were exposed to a continuous stream of auditory and visual stimuli. The stream contained adjacent dependencies between stimulus pairs, with each stimulus in the pair being deterministically associated with another at 100% probability. Our design incorporated both within-modality (unimodal) and across-modality (crossmodal) pairs to compare how effectively people learn these different types of patterns.

For the first two experiments, we deliberately chose synthetic stimuli without semantic meaning: computer-generated fractal images and artificial sounds (see Fig. 1A). This design allowed us to determine whether crossmodal statistical learning could occur at a sensory level rather than through semantic associations. Across all experiments, we assessed statistical learning through two measures: an implicit measure during exposure (online) and an explicit measure after exposure (offline). Experiment 1 sought to validate previous findings by (Walk & Conway, 2016), unimodal and crossmodal pairs were presented in random order during training. In our study we introduced an implicit learning measure, in addition to the explicit judgement used in their original study. Building on this, Experiment 2 investigated whether people’s natural tendency to attentionally group same-modality stimuli together might interfere with learning crossmodal patterns. To test our second hypothesis, we organized the stimuli into distinct blocks by modality combination, allowing participants to anticipate switches between visual and auditory input. This modification was designed to counteract any perceptual bias towards grouping together stimuli of the same modality. For Experiment 3, and to test the third hypothesis, we shifted to using meaningful auditory and visual stimuli to test whether crossmodal statistical learning becomes possible when inputs from different senses can be represented at higher-level, through modality-independent (amodal) representations.

**Figure 1.**
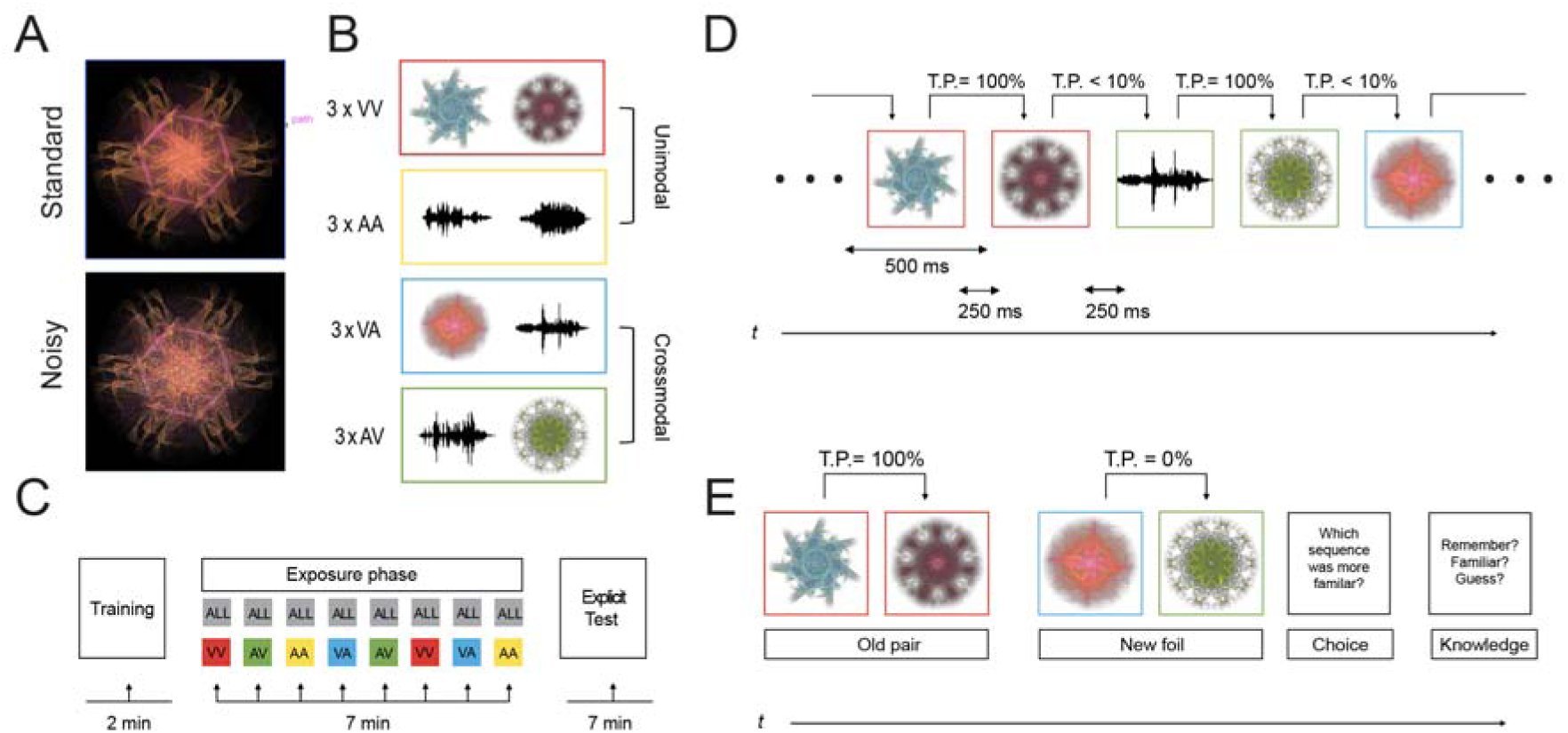
A Example of a standard visual stimulus and its noisy version in the Experiments 1 and 2 B 12 visual and 12 auditory meaningless synthetic (Experiment 1 and 2) or meaningful natural (Experiment 3; not displayed here) stimuli were combined conforming 12 standard pairs (3 x each modality combination). C Each experiment consisted in three parts: First, a training phase in which the participants were familiarized with the task and the different stimuli. Second, an exposure phase with the same number of blocks in all experiments. In Experiment 1 and 3, during the exposure phase, all the modality combinations were randomly intermixed in each block (“ALL” grey square in the top row of the Exposure phase in panel C). In Experiment 2 only one modality combination was presented in each block (depicted with different colors in the bottom row of the Exposure phase in panel C). Third, after the exposure phase, participants performed an explicit recognition test. D Example of a stimuli sequence during the exposure phase in the first experiment (for illustrative purposes, color frames denote standard pairs as represented in B and C). E An example trial of the explicit 2AFC test. The “Old” pair is one of the previously presented pairs and a “New foil” pair is a new stimuli combination that was never presented during the exposure phase. The color frame is added here for illustrative purposes: red for VV, yellow for AA pairs, green for AV and blue for VA pairs.

Foreshadowing our results, we found that when unimodal and crossmodal stimuli transitions are randomly intermixed in during the exposure phase, participants only reliably learned the associations between the unimodal pairs (Seitz et al., 2007; Walk & Conway, 2016). However, when crossmodal transitions were predictable, participants could implicitly learn the association between crossmodal pairs. Finally, we demonstrated that participants learned crossmodal transitional probabilities (even when the upcoming modality was unpredictable) if the sensory inputs in the different modalities are meaningful, validating the hypothesis that statistical learning also occurs between higher amodal stimuli representations, regardless of their input modality.

## Experiment 1

In the first experiment, we investigated whether participants could implicitly learn unimodal and crossmodal statistical regularities between meaningless auditory and visual stimuli. To assess this, we measured implicit statistical learning by recording participants’ reaction times in detecting predictable and unpredictable noisy stimuli during the exposure phase. We then used linear regression models to examine whether detection of predictable noisy stimuli, compared to unpredictable ones, improved with increasing exposure during this phase. After the exposure phase, we assessed whether participants could explicitly recognize the pairs with a two-interval forced choice task.

## Methods

### Participants

Forty-five university students participated in Experiment 1 (40 females, mean age 20.36, range 18-28). None of the participants reported history of auditory problems, and all had normal or corrected-to-normal vision. All participants signed informed consent before the experiment following the guidelines of the Ethics committee of the University of Barcelona. Participants were paid with course credit for their participation. The sample size for this and subsequent experiments was set to a minimum of forty-five participants in consistence with similar studies and considering a medium effect size (Cohen’s d > 0.5) with a power of 95% in the explicit test.

### Stimuli

Visual stimuli consisted of twelve artificially generated images of symmetric round fractals. Images were created in Apophysis 7x, a fractal flame renderization software that uses iterated function systems (Spotworks & Berthoud, 2008). The luminance saliency of the selected stimuli was normalized using histogram equalization applied to the value channel of their HSV (hue, saturation, value) representation. Every image had a diameter of 800 px. And was presented in the middle of a CRT monitor, on a black background.

Auditory stimuli were created using random parametrization of multiple synthesizers from the digital audio workstation FL Studio. Twelve different, unequivocally distinguishable abstract sounds were generated. Stimuli were then edited using the software Audacity: their amplitude was normalized, and their length reduced to 500 ms by cutting and time-stretching. To remove any clicking in the sounds an envelope of 100 ms of attack and 100 ms of release was applied. Auditory stimuli were presented through headphones to reduce interference with any possible environmental noise.

Noisy versions of all stimuli were created to have a measure of implicit learning during the exposure phase. In the visual modality, a layer of upscaled RGB white Gaussian noise with the same shape as the stimuli was added to each stimulus. The noise was upscaled to get an additional pixelated effect (Fig. 1A). In the auditory modality, white Gaussian noise with the same amplitude as the stimuli was added to the sounds. Visual and auditory noisy stimuli were designed so that they could be easily discriminated from the standard stimuli while still sharing enough characteristics with their corresponding standard stimulus.

### Procedure

The participants seated in a dimly light, sound attenuated room, facing a computer screen. The viewing distance from the participants to the monitor was approximately 50 cm. Before the experiment, participants were informed they would watch and listen to a series of images and sounds that could either be standard or noisy. To ensure that the participants understood the difference between standard stimuli and noisy stimuli, they were presented with an example of a stimulus per modality and their corresponding noisy version before starting. Instructions were given to the participants both orally by the experimenter and in written form printed on the screen.

The experiment consisted of three phases (Fig. 1C): a *training phase*, in which participants got familiar with the stimuli and the task; an *exposure phase*, in which they were exposed to the stimuli paired following the statistical regularities; and an explicit test phase, in which participants were tested regarding their knowledge acquired from such regularities. Each phase was separated by a short break of 1 minute.

During the training and exposure phases, the participants’ task consisted of detecting noisy stimuli by pressing the spacebar key whenever they saw or hear one. The task was orthogonal to the hidden structure of pairs that randomly appeared during the stream. Participants were instructed to direct their gaze at a white fixation cross at the center of the monitor. Each visual or auditory stimulus was presented for a duration of 500 ms and they were separated by a 250 ms interval.

In the *training phase*, all the stimuli were presented to the participant in random order, without any paired structure. Noisy stimuli appeared at a higher rate than during the exposure phase (35% of the stimuli), because the objective of this phase was to teach participants to discriminate between standard and noisy stimuli. The training phase lasted for one minute, but it was repeated until the participant was able to detect at least the 80% of the noisy stimuli correctly. This way we ensured that the participant would perform the noisy stimuli detection task correctly.

The *exposure phase* had the objective to expose the participants to learn the transitional probabilities and measure their implicit learning of the pairs. In this phase, visual and auditory stimuli (twenty-four in total) were pseudo-randomly grouped into twelve pairs conforming four different modality combinations (Fig. 1B): auditory to auditory (AA), visual to visual (VV), auditory to visual (AV) and visual to auditory (VA), with three pairs for each modality combination. All four conditions were randomly sorted conforming one single stream (Fig. 1D). Transitional probabilities between the stimuli within a pair was set at 100%, while the transitional probabilities between stimuli of different pairs was set at 10%. Each pair was presented 110 times during exposure leading to a total of 1320 pair presentations at the end of the exposure phase. Pairs were therefore presented in random order with the only restriction that the same pair could not repeat after itself. Each stimulus appeared in its noisy version 10 times, out of a total of 110. Noisy appearances were randomly scattered across the exposure phase and they appeared half of the times on the first item of the pair (leading stimulus) or on the second item of the pair (trailing stimulus).

This part of the experiment was divided into eight blocks, each of them lasting for approximately six and a half minutes. Between blocks, participants had the opportunity to rest and then resume the experiment by pressing a key whenever they wanted. In order to maintain the participants engaged with the task, after each block, participants received feedback about the percentage of noisy stimuli detected. To constrain the temporal window between noisy stimuli and keypresses and calculate the proportion of correct noisy stimuli detection, we considered a correct detection whenever a keypress occurred no later than 2 seconds after the noisy stimulus’ onset.

Following the exposure phase, we informed the participants for the first time that they would need to complete an additional task. In this *explicit test phase*, we explained that the visual and auditory stimuli they encountered during the exposure phase were presented conforming pairs. To evaluate whether the participants had explicit knowledge of these pairs, we administered a two-alternative forced choice (2AFC) task. In each trial, participants were sequentially shown an old pair and a new foil pair, with the deviant item equally often inserted in either leading or trailing positions. The old pair had been presented during the exposure phase, while the foil pair violated the statistical regularities of the exposure phase.

For each condition, the stimuli included three old pairs from the learning phase (e.g., for visual pairs: V1-V4, V2-V5, V3-V6), and three new foil pairs of the same modality. These foils were created by randomly pairing the leading stimuli of each standard pair with a trailing stimulus from another pair (e.g., V1-V5, V2-V6, V3-V4). Importantly, these foils were new to the participants and were matched in frequency of occurrence for each stimulus, ensuring that participants could not rely on differences in stimulus frequency to solve the task. This procedure was repeated for each modality, resulting in a total of 24 pairs (12 old and 12 foil). Each old pair was presented against three foil pairs of the same modality (e.g., V1-V4 vs. V1-V5, V1-V4 vs. V2-V6, V1-V4 vs. V3-V4), yielding a total of 36 test trials.

After viewing both old and new foil pairs in sequence, participants selected, using the left and right arrow keys, which pair felt more familiar to them (left for the first pair, right for the second). They were also asked to report the confidence of their decision by pressing ‘Z’ if they “remembered the sequence,” ‘X’ if “the sequence was more familiar,” or ‘C’ if they were “guessing.” These responses correspond to three confidence levels: Remember, Know, and Guess, respectively. We treated the Remember and Know responses as indicators of explicit knowledge, accessed through either recall or familiarity. The test was self-paced: participants pressed a key to initiate each pair and had no time limit for responding.

### Analyses

To evaluate whether participants could implicitly learn the statistical regularities, we focused our analysis on reaction times (RTs). We included only RTs longer than 200 ms and shorter than 2 seconds relative to the noisy stimulus onset. To reduce the positive skewness typical of RT distributions, we normalized them by calculating the reciprocal of the latencies (Whelan, 2008), where higher 1/RT values indicate faster responses.

To assess learning, we examined the impact of noisy stimulus position (leading vs. trailing within each pair) on detection speed using a linear mixed-effects model (LMM). LMMs offer greater statistical power than classical ANOVA analyses because they can account for variability in RTs between participants or stimuli as random effects. Specifically, we quantified implicit statistical learning by modelling RTs as a function of the noisy stimulus position within the pair (i.e., the “learning” regressor).

Based on prior research (Barakat et al., 2013; Richter & de Lange, 2019; Yan et al., 2021; Zhou et al., 2019), we predicted that participants would demonstrate implicit learning by using the leading stimulus to anticipate the trailing stimulus in each pair. This anticipation must lead to faster RTs for detecting trailing compared to leading noisy stimulus, reflecting implicit learning throughout the exposure phase (Fig. 2, middle panel).

**Figure 2.**
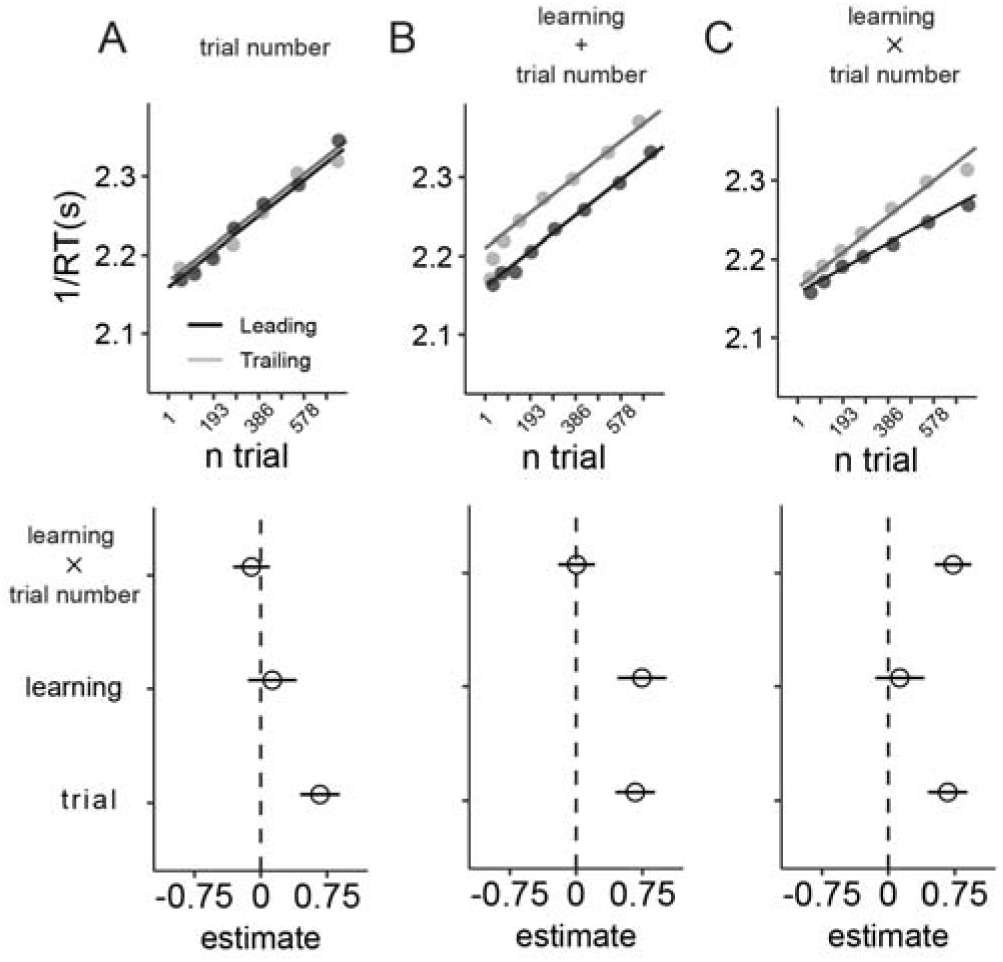
Implicit Statistical Learning Model Regressors Predictions. The upper rows represent artificially generated linear regression model predictions for different possible experimental outcomes, while the bottom rows depict the corresponding parameter estimates of the models described above. Interpretation of parameter estimates: The ‘intercept’ of the model represents the estimate of 1/RT at the beginning of the experiment (baseline 1/RT prior to any learning). The ‘trial’ regressor estimate indicates the modulation of 1/RT as a function of the number of trials. The ‘learning’ regressor represents the average difference in 1/RT between trailing and leading noisy stimuli. Finally, the learning rate is computed as the linear interaction between the number of trials and the learning regressor. A. Shows the parameter combination for an outcome where there is no statistical learning, with performance improving as the experiment progresses. B. Shows the parameter combination for an outcome where participants successfully learned the statistical regularities. A significantly positive value for the learning regressor indicates that participants promptly learned the statistical associations, as performance differences appear from the beginning of the experiment. C. Shows a positive learning rate, indicating that participants increasingly utilized the predictive association between leading and trailing items, with the learning gradually unfolding over time. The grey and black points in the upper row represent the artificially generated underlying 1/RT averages of trailing and leading stimuli, respectively, across trials.

To assess continuous and sustained implicit learning of transition probabilities throughout the experiment, we used the “trial” number of each stimulus presentation as a continuous regressor in relation to the learning regressor. The trial number reflects the number of times a specific modality combination was presented during the implicit phase. For instance, a trial number of 600 for the crossmodal conditions indicates that participants had encountered 600 crossmodal pairs in the exposure phase. Importantly, the interaction between the “learning” and “trial” regressors (Fig. 2C) represents the learning rate, which captures sustained variations in learning across multiple trials. To improve the mixed model’s convergence, we rescaled trial number regressor to z-scores.

To test for potential differences in implicit learning between unimodal and crossmodal transitions, in our main analyses we grouped auditory and visual conditions together (unimodal transitions) and audiovisual and visuo-auditory conditions together (crossmodal transitions) (see Table 1; Eq. 1). To account for variability due to the modality of the noisy stimulus, we included “modality” (visual/auditory) as a regressor of no interest and as an additional random-effect term, addressing modality-specific differences it might cause on RT. We also included random error effects, allowing the intercept to vary independently for each participant and stimulus identity. In addition, we performed exploratory analyses to test whether statistical learning within the unimodal and crossmodal conditions was driven by the auditory, the visual noisy stimulus or both.

**Table 1:** Linear Mixed Models for Quantifying Implicit and Explicit Statistical Learning Effects. Eq.1 depicts the full Linear Mixed Model (LMM) model used to quantify implicit learning from speediness measurements (1/RT). We compared our base model, which includes a trial regressor and its interaction with the modality transition regressor, against other extended versions of the model including a main effect [+] or an interaction [*] with the learning regressor. Eq.2 represents the Generalized Linear Mixed Model (GLMM) utilized to predict the probability of correct responses based on the confidence level. Random effects of the mixed models include (modality|participant) and (1|stimulus).

|  | Dependent variable | Regressors | Equation ID |
| --- | --- | --- | --- |
| Implicit statistical learning effects | 1/RT | ~ modality + trial * modality transition [+ or *]<br>learning (noisy stimulus position) + Random effects | Eq. 1 |
| Explicit statistical learning effects | P(Correct) | ~ confidence level + Random effects | Eq. 2 |

The models were optimized using the bobyca algorithm from the lme4 package (Baayen et al., 2008; Bates et al., 2014) in R (version 3.6.2, R Foundation for Statistical Computing). We used post-hoc likelihood-ratio (χ^2^) model comparisons to assess the predictive power and significance of all main effects and interactions. A reduced model, containing only trial number and modality transition as fixed effects, was compared to increasingly complex models, including the learning regressor. After multiple comparisons, we identified a model that best explained the data, reporting the obtained χ^2^ against the immediately less complex model. We then assessed the significance of each fixed effect estimate by calculating the Wald (z) statistic in the full model. For additional support of a null hypothesis, Bayes-factor (BF) estimates from model comparisons showing null effects are reported.

Figures in this manuscript display regressor coefficients and confidence intervals from the most complex model, which includes all main effects and interactions, allowing for direct comparison across experimental conditions.

To test whether participants could explicitly recognize the associations between the leading and trailing stimuli, we modelled the responses of the 2AFC explicit learning test using GLMM (family binomial logit) implemented in lme4. As this task consisted of a 2AFC, chance level is equal to a 50% of correct responses (logistic regression intercept = 0). Similarly to the implicit learning analyses, we modelled unimodal and crossmodal modality transitions conditions separately. This approach was primarily motivated by the low number of trial repetitions for each modality combination. If the intercept of the model is significantly larger than 0, it means that participants can discriminate the old pairs from the foils above chance level (>50%). Moreover, to determine whether the participants guessed or explicitly recognized the old pairs, we also modelled the level of confidence reported in the decision (Table 1; Eq.2). We only report the results and statistics of the post hoc model comparisons in the main results section. Again, the reported χ2 values reflect the contrast of the best model against the immediately simpler contrasted model. The stimuli and code used to run the experiments, and reproduce the analyses and figures is available at https://osf.io/tcxe9/ (Pérez-Bellido, 2023).

## Results

### Exposure phase: implicit learning for unimodal pairs only

The high accuracy of the participants in noise detection (Fig. 3A) indicated thar participants were paying attention to both modalities (group average: P(Hits)_v_ = 0.99 and P(Hits)_A_ = 0.85). A repeated measures ANOVA showed that participants detected more noisy stimuli in the visual compared to the auditory conditions (F(1, 63) = 52.2, P < 0.01), but their detection performance did not change as a function of modality transition (F(1, 63) = 0.063, P = 0.8).

**Figure 3.**
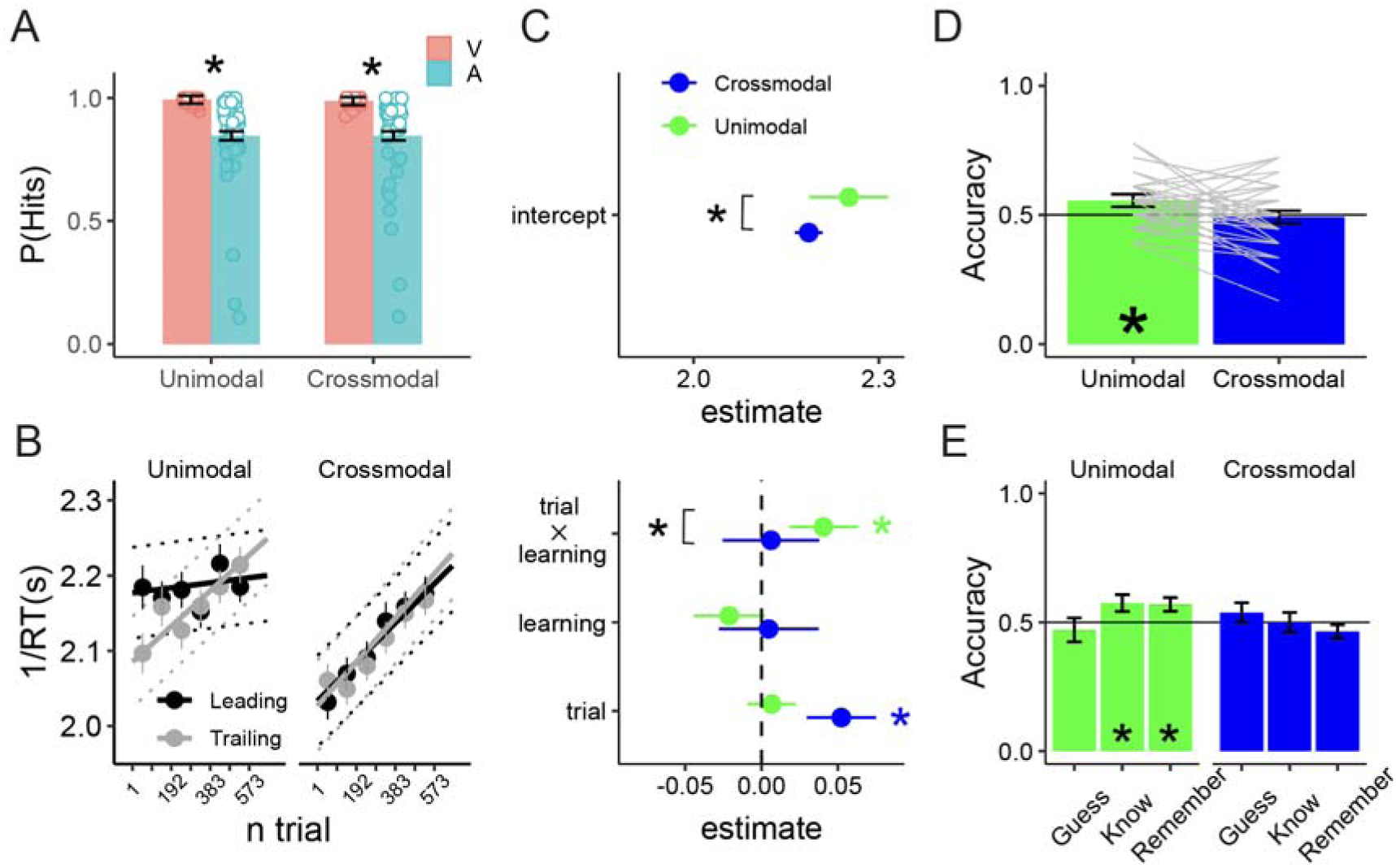
Performance during the exposure and explicit learning phases in Experiment 1: A. Group-average speediness (1/RT) for the unimodal and crossmodal pairs, separated by stimulus modality (V, A) and modality transition (unimodal, crossmodal). B. Full model predictions for the unimodal and crossmodal pairs. Thin lines represent the confident intervals of the model. Filled points with error-bars represented group-level average values at different trials bins and their corresponding standard error. C. Parameter estimates of the full model depicted in B. D. Explicit test accuracy in discriminating old from new foils in the unimodal and crossmodal conditions. E. Analysis of participants accuracy as a function of the confidence. Confidence intervals (error bars) are calculated as the parameter estimate standard error multiplied by 1.96. Based on model parameter contrasts, black asterisks indicate significant differences between conditions estimates, while colored asterisks indicate significant modulations of a given parameter with respect to 0.

We used linear mixed-effects models (LMMs) and conducted post hoc likelihood-ratio (χ^2^) model comparisons to investigate the impact of trial number and learning regressors on participants’ response times for detecting noisy stimuli (as illustrated in Fig. 3B and C). Our analysis showed that the full model, which included a triple interaction between trial number, modality transition, and learning, provided the best explanation for the data (χ^2^ = 14, df = 3, P < 0.01). A closer look at the regressors fitted to the unimodal trials showed that participants’ noise detection speed did not differ between leading and trailing noisy stimuli at the beginning of the experiment (χ^2^ = 1.2, df = 1, P = 0.2; BF = 0.02), as indicated by the non-significant difference between the leading and trailing intercepts (learning: z = -1.7, P = 0.07). However, as the experiment progressed and the number of trials increased, participants became faster in detecting trailing noisy stimuli compared to leading ones (learning rate: z = 3.46, P < 0.01), consistent with an implicit learning effect (Fig. 2C). We replicated similar effects for the unimodal auditory (learning rate: z = 2.3, P < 0.05) and visual (learning rate: z = 3.2, P < 0.005) noise, demonstrating that unimodal statistical learning was not primarily driven by either modality. In contrast, we did not observe a similar learning effect in the crossmodal pairs (learning rate: z = 0.3, P = 0.78; Model comparison χ^2^ = 0.01, df = 2, P = 0.95, BF < 0.01). Overall, these findings suggest that participants gradually acquired and utilized the stimulus associations to improve noise discrimination only for the unimodal pairs.

Additional analyses revealed that participants’ RTs on average were faster for visual noise compared to auditory noise (modality: z = -4.1, P < 0.001), suggesting that detecting auditory noise is more effortful than detecting visual noise, and improved with the number of trials in the crossmodal pairs (trial: z = 7.4, P < 0.001). Moreover, differences between unimodal and crossmodal intercepts indicated that participants, on average, were slower in detecting noise during crossmodal modality transitions (modality transition: z = -5.6, P < 0.001), consistent with the modality switching cost observed in the proportion of hits for noisy stimuli detection.

### Explicit test phase: explicit learning of unimodal transitional probabilities

The GLMM analyses of correct response proportions in the explicit learning test phase revealed that participants could recognize unimodal old pairs above chance level (intercept: z = 3.61, P < 0.001; χ2 = 10.02, df = 1, P < 0.005). However, we found no evidence of old pair recognition for crossmodal conditions (χ2 = 2.93, df = 1, P = 0.23; BF = 0.3; Fig. 3D). Additionally, the confidence level regressor only improved the model’s fit in the unimodal conditions (χ2 = 6.1, df = 2, P < 0.05). Specifically, participants showed a higher proportion of correct responses in both the “know” (z = 2.65, P < 0.005) and “remember” (z = 2.18, P < 0.05) conditions compared to the “guess” condition (Fig. 3E). These results suggest that participants could only explicitly recognize the unimodal pairs.

In summary, this experiment shows evidence of unimodal implicit and explicit statistical learning. However, the results regarding crossmodal statistical learning align with previous studies showing a modality-specific limitation that prevents the extraction of regularities between modalities at the sensory level.

## Experiment 2

In our first experiment, we observed that participants were able to learn and use unimodal transitional probabilities effectively to discriminate the noisy stimuli, while crossmodal transitional probabilities were not learned. These findings support the notion that statistical learning is a modality-specific process that operates within each sensory modality (Frost et al., 2015). However, it is important to note that in Experiment 1, stimuli were presented in a continuous stream, where unimodal and crossmodal transitions were randomly intermixed. This random presentation may have hindered the learning of crossmodal pairs due to the perceptual costs associated with switching between modalities, as previously demonstrated in multisensory research (Gondan et al., 2004; Otto & Mamassian, 2010; Spence et al., 2001). To reduce the potential impact of this attentional cost on statistical learning, we designed our second experiment to include eight different experimental blocks, each containing a specific combination of visual-visual (VV), auditory-auditory (AA), audio-visual (AV), and visual-audio (VA) stimuli (Fig. 1C). By doing so, participants could anticipate the modality of the incoming stimuli in the crossmodal blocks (e.g., AVAVA or VAVAV), thus eliminating the possibility of a bias to perceptually group consecutive visual or auditory stimuli together.

## Methods

### Participants

Sixty-three university students participated in Experiment 2 (59 females, 4 males, mean age 20.36, range 18-28). Participant recruitment criteria and stimuli were identical to those in the previous experiment. Due to an error in participant recruitment, we collected 13 more participants than our target (45 + 5 possible dropouts). Since eliminating the last 13 participants from the final sample did not change the results, we decided to maintain them in the analyses.

### Procedure

The experimental procedure was almost identical to Experiment 1. However, here there was only one modality combination in each experimental block, being either VV, AV, AA, or VA (Fig. 1C). The order of each modality block was randomized for each participant, and two blocks of the same modality combination could not come one after the other. We kept the transitional probabilities between the stimuli within a pair at 100%. Importantly, because we kept the stimuli identical to Experiment 1 varying only the stimuli presentation, this implied a smaller number of possible pairs within each block (only 3 pairs, 6 stimuli). Therefore, the transitional probabilities between pairs was 0.5 (more than ×4 higher than in Experiment 1), potentially rendering the statistical learning of the associated pairs more challenging compared to the previous experiment.

## Results

### Implicit learning of crossmodal transitional probabilities only

During the exposure phase (Fig. 4A), participants successfully detected most of the visual and auditory noisy stimuli (P(Hits)_V_ = 0.99 and P(Hits)_A_= 0.88). A repeated measures ANOVA showed that participants detected more noisy stimuli in the visual compared to the auditory conditions (F(1, 63) = 52.2, P < 0.01), but their detection performance did not change as a function of modality transition (F(1, 63) = 0.063, P = 0.8).

**Figure 4.**
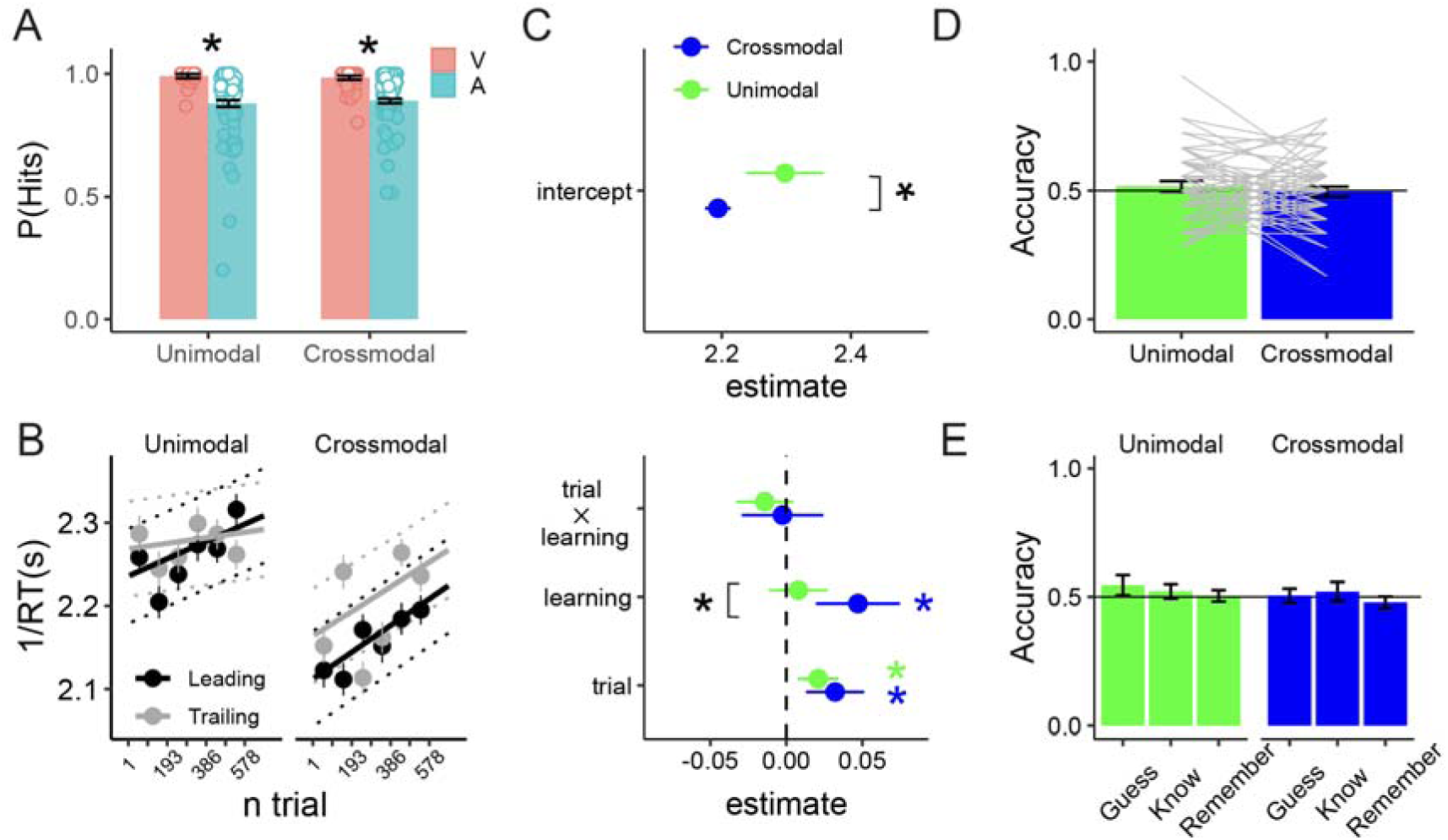
Performance during the exposure and explicit test phases in Experiment 2: A. Group-average speediness (1/RT) for the unimodal and crossmodal pairs, separated by stimulus modality (V, A) and modality transition. B. Full model predictions for the unimodal and crossmodal pairs. Thin lines represent the confident intervals of the model. Filled points with error-bars represented group-level average values at different trials bins and their corresponding standard error. C. Parameter estimates of the full model depicted in B. D. Explicit test accuracy in discriminating old from new foil pairs in the unimodal and crossmodal conditions. E. Analysis of participants accuracy as a function of the confidence. Confidence intervals (error bars) are calculated as the parameter estimate standard error multiplied by 1.96. Based on model parameter contrasts, black asterisks indicate significant differences between conditions estimates, while colored asterisks indicate significant modulations of a given parameter with respect to 0.

To investigate whether participants learned the item associations, we used a linear mixed model (LMM) to analyze 1/RTs as a function of trial number (full model in Fig. 4B and C). The best-fitting model included trial number, type of modality transition, and learning as fixed effects (χ^2^ = 10.13, df = 3, P < 0.02). A closer look at the parameter’s estimates showed that the learning effect was driven by the crossmodal conditions, as on average, participants were faster in detecting crossmodal trailing than leading noise (learning: z = 4.6, P < 0.001). This difference did not continue increasing throughout the number of trials (χ^2^ = 0.16, df = 2, P = 0.9, BF < 0.01), as the learning rate parameter was not significantly different from 0 (learning rate: z = -0.4, P = 0.68). That is, the learning of crossmodal statistical regularities reaches its maximum in a relatively small number of trials and does not continue evolving with additional exposure. We replicated significant positive differences in the learning parameter when analysing the auditory (learning_A_: z = 3.7, P < 0.001) and visual (learning_V_: z = 3.2, P < 0.005) crossmodal conditions separately, demonstrating that crossmodal statistical learning was not driven by either of the two possible crossmodal modality combinations.

Our complementary analyses on the unimodal trials did not provide evidence that participants learned the unimodal transitional probabilities, as the RTs in detecting leading and trailing noisy stimuli were not significantly different (χ^2^ = 1.4, df = 1, P = 0.23, BF = 0.05).

This rapid statistical learning (Fig. 2B), which contrasts with Experiment 1 where implicit learning developed gradually, may be attributed to differences in the experimental design. In this new experiment, each modality combination pair was repeatedly presented within its experimental block (e.g. AV block), whereas in Experiment 1, modality combinations appeared throughout the entire experiment. Frequent exposure to the same pairs over a few trials may enhance the learning of statistical dependencies.

Additionally, the analyses revealed that on average, participants’ reaction times improved with the number of trials for both the noisy stimuli in unimodal and crossmodal pairs (trial: z = 6, P < 0.001). This sustained and general RTs improvement can be attributed to a practice effect. Participants also showed faster responses to visual compared to auditory noise (modality: z = -2.14, P = 0.04) suggesting that it was easier to detect visual noise. Finally, based on the significant differences between the unimodal and crossmodal model intercepts, we replicated that, on average, participants were slower in detecting a noisy stimulus preceded by a stimulus in a different modality (modality transition: z = -10.1, P < 0.001). These results indicate that in Experiment 2, despite participants were able to anticipate modality alternations due to the task structure, they still incurred in a cost associated with modality switches.

### No evidence of explicit statistical learning

GLMM analyses on the proportion of correct responses showed that participants could not explicitly recognize the unimodal (χ2 = 0.93, df = 1, P = 0.3, BF = 0.04) nor the crossmodal pairs (χ2 = 0.1, df = 1, P = 0.7, BF = 0.03). Overall, the results of this second experiment show that participants were able to benefit for the predictable alternation between modalities to learn crossmodal transitional probabilities and use them to improve their detection of trailing noisy stimuli but this was not reflected in the explicit measures of learning. The unimodal conditions were more penalized by the higher transitional probabilities at boundaries between pairs in comparison to the first experiment than the crossmodal pairs. These findings suggest that crossmodal statistical learning occurs when modality transitions are predictable. Additionally, our results indicate that crossmodal statistical learning is primarily evident when assessed through implicit measures. In contrast, we also demonstrate that unimodal statistical learning can be observed through both implicit and explicit indices, provided there is a sufficient difference in the transitional probabilities between paired and non-paired items.

## Experiment 3

In Experiment 3, we tested the prediction that both unimodal and crossmodal regularities can be learned if the stimuli contain semantic information and are therefore not tight to the sensory level characteristics. Consistent with the hypothesis proposed by Frost et al., (2015), we predicted that if statistical learning can occur between high-level stimuli representations (Brady & Oliva, 2008), participants should be able to learn crossmodal contingencies between meaningful crossmodal pairs. To test this prediction, we used visual and auditory stimuli with meaningful content to facilitate the formation of sensory-independent stimulus relations.

## Methods

### Participants

Fifty university students participated in Experiment 3 (7 males, mean age 21.9, range 18-45). Participant’s recruitment criteria and the stimuli were identical to the previous experiments.

### Procedure

Similar to the first experiment, we randomly intermixed visual, auditory, audiovisual, and visuo-auditory pairs in each experimental block while maintaining the transitional probabilities between stimuli within a pair at 100% and the transitional probabilities between pairs at 10%. However, unlike the previous experiments, we used meaningful stimuli to represent visual and auditory objects. The visual images were composed of the same number of visual and auditory items than Experiments 1 and 2. Six animate objects (e.g., crocodile) and 6 inanimate objects (e.g., piano) were overlaid on a black background. Similarly, auditory stimuli consisted of 500 ms clips corresponding to 6 animate (e.g., dog barking) and 6 inanimate objects (e.g., phone ringing). All stimuli were freely downloaded from the internet and pre-processed in the same manner as the fractals and synthetic stimuli used in the previous experiments to generate the noisy version of the stimuli. Prior to the beginning of the experiment, the experimenter presented all the stimuli to the participants one by one to ensure that they understood the semantic referent of each image and sound. We found that all the stimuli referents could be correctly identified by all the participants.

## Results

### No evidence of implicit statistical learning

During the exposure phase, participants accurately detected both the visual and auditory noisy stimuli, with group averages of P(Hits)_V_ = 0.97 and P(Hits)_A_ = 0.84 (Fig. 5A). To explore how participants’ noise detection changed as a function of modality and modality transition, we conducted a repeated measures ANOVA. This analysis revealed that participants detected more noisy stimuli in the visual than auditory modalities (F(1,48) = 75.6, P < 0.01) and more noisy stimuli in the crossmodal than unimodal pairs (F(1,48) = 75.6, P < 0.01). We also observed a significant interaction between modality and modality transition (F(1,48) = 15.5, P < 0.01), indicating that auditory noise was better detected in the crossmodal than unimodal pairs

**Figure 5.**
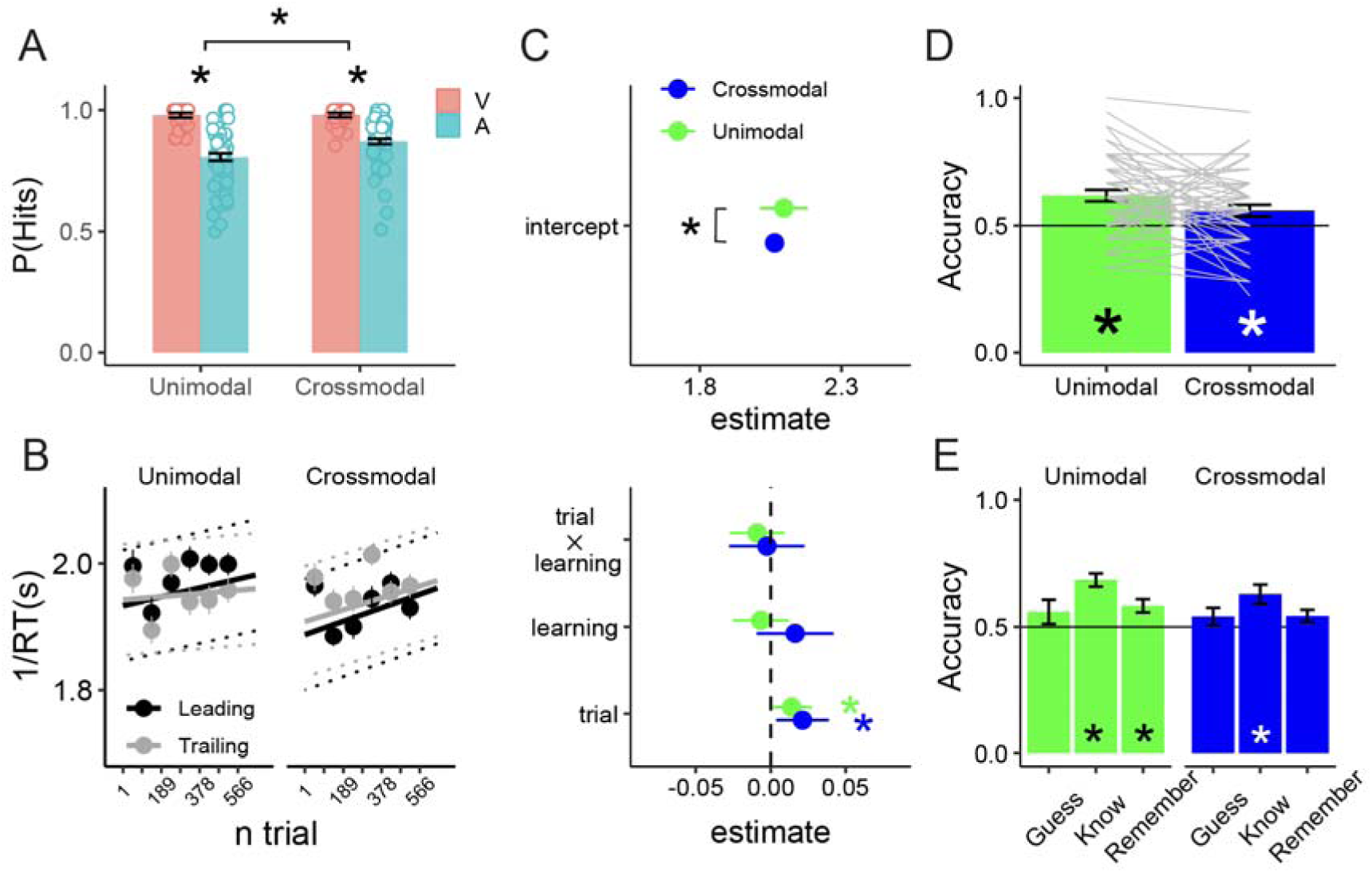
Performance during the exposure and explicit learning phases in Experiment 3: A. Group-average speed (1/RT) for the unimodal and crossmodal pairs, separated by stimulus modality (V, A) and modality transition. B. Full model predictions for the unimodal and crossmodal pairs with confident intervals. Thin lines represent the confidence intervals of the model. Filled points with error-bars represented group-level average values at different trials bins and their corresponding standard error. C. Parameter estimates of the full model depicted in B. D. Explicit test accuracy in discriminating old from new foil pairs in the unimodal and crossmodal conditions. E. Analysis of participants accuracy as a function of the confidence. Confidence intervals (error bars) are calculated as the parameter estimate standard error multiplied by 1.96. Based on model parameter contrasts, black asterisks indicate significant differences between conditions estimates, while colored asterisks indicate significant modulations of a given parameter with respect to 0.

We fitted a LMM on reaction times to determine whether participants learned pair associations implicitly (Fig. 5B and C). Our analysis revealed that the best model for explaining participants’ responses only included trial and modality transition as fixed-effect regressors (χ2 = 22.8, df = 1, P < 0.01). We did not find evidence of implicit statistical learning, as including the learning regressor did not improve the model fit (χ2 = 0.58, df = 1, P = 0.44, BF < 0.01). Further investigation of the regression estimates showed that participants’ performance improved with increased practice in the unimodal and crossmodal conditions (trial: z = 1.91, P = 0.04), indicating a practice effect. Participants also showed faster responses to visual compared to auditory noisy stimuli (modality: z = -5.3, P < 0.001). Furthermore, in line with previous analyses, the differences between unimodal and crossmodal intercepts indicated that participants were generally slower at detecting noise in stimuli when these were in crossmodal than unimodal pairs(modality transition: z = -3.4, P < 0.001), reflecting a modality switching cost. Overall, these findings did not provide empirical evidence of statistical learning through the implicit measure of noise detection in either modality or during modality transitions.

### Explicit learning of unimodal and crossmodal pairs

GLMM analyses on the proportion of correct responses in the explicit learning test phase reveal that participants did recognize the unimodal old pairs above chance level (χ2 = 20.9, df = 1, P < 0.001; Fig. 5D). Interestingly, we also found that participants explicitly recognized the crossmodal old pairs (χ2 = 6.3, df = 1, P < 0.02). The confidence level regressor significantly improved the model’s fit in both the unimodal (χ2 = 15.8, df = 2, P < 0.01) and the crossmodal conditions (χ2 = 7.8, df = 2, P < 0.03). Participants reported a larger proportion of correct responses in the “know” confidence level compared to the “remember” (know vs remember; unimodal: z = 3.4, P < 0.005; crossmodal: z = 2.7, P < 0.01) and “guess” (know vs guess; unimodal: z = 3.2, P < 0.005; crossmodal: z = 2.2, P < 0.03) levels highlighting the explicit nature of the observed learning (Fig. 5D).

These results indicate that participants can recognize both unimodal and crossmodal old pairs when they contain meaningful information. Moreover, participants showed a degree of awareness in their learning, as evidenced by greater accuracy in decisions rated with a confidence level of know.

## Discussion

Previous studies have failed in showing crossmodal interactions in statistical learning (Conway & Christiansen, 2006; Seitz et al., 2007; Walk & Conway, 2016). This has led to propose that, although statistical learning relies on domain-general computational principles, the learning of statistical dependencies between multimodal low-level stimulus features (e.g. tones, shapes, vibrations) is constrained by the representational characteristics of its correspondent sensory modality (Frost et al., 2015). Here, in three separate experiments we investigated different explanations that could have impeded previous studies to detect crossmodal statistical learning.

In our first experiment, we replicated previous studies that demonstrated significant statistical learning between unimodal stimuli, but null statistical learning between crossmodal stimuli (Walk & Conway, 2016). Our results provided evidence of statistical learning through both explicit task responses and the use of an implicit measure. Specifically, we found that while reaction times towards leading and trailing noise detection were not significantly different at the beginning of the implicit training phase, as the experiment progressed, participants became faster in detecting the noisy stimuli in trailing compared to the leading positions in unimodal pairs, indicating implicit learning and exploitation of the unimodal associations. Crossmodal learning was not observed even when learning was measured with implicit measures during the learning phase.

One possible explanation for the null crossmodal statistical learning in Experiment 1 is that unimodal and crossmodal stimuli pairs were randomly intermixed during the exposure phase. This mixing may have introduced modality switching costs, potentially biasing participants’ attention toward grouping inputs from the same sensory modality together. This attentional bias might facilitate the learning of unimodal associations but impair the learning of crossmodal contingencies, as previous research has highlighted the crucial role of attention in both unimodal (Treisman & Gelade, 1980) and crossmodal (Talsma et al., 2010) information binding, as well as in statistical learning processes (López-Barroso et al., 2016; Richter & de Lange, 2019; Toro et al., 2005). To address this potential confound, in Experiment 2 we tested whether participants could learn crossmodal associations more effectively when stimuli were paired in modality-predictable transitions, thereby eliminating the attentional bias toward grouping stimuli from the same modality. Notably, the reaction time analysis in Experiment 2 revealed that participants were able to implicitly learn and exploit the crossmodal associations, contrasting with their inability to learn the unimodal associations. While the latter were hindered by the increased transitional probabilities at the boundaries between pairs in Experiment 2, rising from 10% to 50% with respect to Experiment 1, the alternation between modalities might have helped to mitigate this limitation in crossmodal pairs. In this condition, participants could use the modalities alternation as a cue to know which modality was the leading item once one pair was accurately segmented. This could have facilitated the learning process even in the face of increased transitional probabilities between items. Alternatively, it is possible that different cognitive mechanisms are involved in learning unimodal versus crossmodal associations, leading to differential effects of transitional probabilities on learning. However, further research is needed to fully understand the underlying mechanisms behind these findings. In any event, the results of Experiment 2 are consistent with our a priori hypothesis, which posited an attentional bias to group stimuli from the same modality together. Such a bias might hinder the capacity to learn the associations between inputs delivered through different sensory modalities when these are not temporally segmented as pairs. In line with this idea, previous research (Kok et al., 2012) using audiovisual associations between low-level stimuli (i.e., frequency tones and grating orientations), but presented in temporally discernible pairs and always and with modality associations occuring always in the same order, has already shown learning of the crossmodal correspondences at both the behavioral and neural levels. In summary, the results of this second experiment highlight the importance of attention as a modulatory factor of statistical learning (Orpella et al., 2021; Perruchet & Pacton, 2006), here guided by the predictability of modality alternations, facilitating crossmodal learning when stimuli are processed only at the sensory level.

Finally, in a third experiment, we tested whether participants could learn crossmodal associations using meaningful visual and auditory stimuli. Drawing from findings demonstrating that humans can learn category-level statistical regularities (Brady & Oliva, 2008; Yan et al., 2023), we aimed to show that humans can also learn transitional probabilities between multisensory inputs that can be encoded in an amodal format, detached from their sensory referents. In line with this hypothesis, in Experiment 3, using an implicit learning task, similar to the one used in Experiment 1, participants were able to learn both the unimodal and the crossmodal associations explicitly. Our results are consistent with previous studies (Cunillera, Laine, et al., 2010) that demonstrate that the semantic processing of the stimuli may facilitate the explicit recognition of the associated pairs in both unimodal and crossmodal conditions.

The inclusion of implicit and explicit measures of statistical learning was essential in this study to be able to observe whether crossmodal SL took place. This was particularly important, in Experiments 2 and 3 where dissociations between those measures were observed. Previous studies have documented the absence of a correlation between those measures (Forscher et al., 2019). For this reason, the inclusion of both is critical in all SL studies since different mechanisms may underlie the two types of learning and/or these measures have different sensitivity to learning. Nevertheless, whereas implicit learning can occur without explicit learning, as we observed in Experiment 2, the opposite is unlikely (Batterink et al., 2015; Conway, 2020). Therefore, we believe that a null effect in implicit learning in Experiment 3 does not necessary mean that there was not implicit learning. Although we do not have a priori explanation for this null result, we speculate that it is possible that the online measure that we used to index implicit learning was not optimal in capturing the learning when using semantic stimuli. Whereas the processing of meaningless stimuli in Experiments 1 and 2 may favor the encoding of sensory differences (i.e., sensory noise), the processing of meaningful stimuli may prioritize the neural encoding of high-level referents of the presented inputs, reducing the impact of statistical learning on the processing of the low-level sensory characteristics of the inputs (Hochstein & Ahissar, 2002). Perhaps using a task in which participants must perform a semantic task would help reveal implicit statistical learning. Indeed, the fact that the stimuli had a semantic content but the task that used to measure implicit learning required focusing at the perceptual level made the detection task more difficult, as reflected by the increased reaction time in Experiment 3 compared to Experiments 1 and 2. This may have impeded to observe learning in the implicit measure due to a reduced sensitivity at such high reaction times.

Our behavioral results do not allow us to draw definitive conclusions about the neural mechanisms involved in crossmodal statistical learning. However, they do support the idea that statistical learning can occur at multiple levels of representation, encompassing both sensory and amodal information. The key difference lies in the fact that while the learning of low-level associations is constrained by perceptual phenomena (i.e., grouping biases), learning of semantic associations bypasses these limitations. This aligns with neural evidence from intracranial recordings (Henin et al., 2021), which shows that both supramodal and modality-specific brain structures are involved in statistical learning across sensory modalities. While sensory areas encode transitional probabilities specific to either auditory or visual statistical learning, the hippocampus and higher-order brain regions—which process the internal structure related to the order and identity of elements—are shared across modalities. The existence of shared statistical learning mechanisms across sensory modalities has also been characterized behaviorally. Sabio-Albert et al. (2025) demonstrated that accuracy in one modality was influenced not only by the predictability of this modality but also by the predictability of an ignored modality. These interactive effects reveal an interdependence between the visual and the auditory statistical learning mechanisms.

In summary, this study demonstrates that humans can indeed learn crossmodal statistical regularities. It identifies several factors that may be determinant for this learning, including the presence of within-modality attentional biases and the level of representation of sensory inputs. Our results contribute to expand the view of statistical learning suggesting that learning of statistical regularities between low-level stimuli features relies on hard-wired learning computations occurring in their respective sensory cortex. Specifically, we show that visual and auditory synthetic stimuli, which are challenging to represent in an amodal format (e.g., semantic representations), can still be bound together, albeit with perceptual limitations that can be overcome through attention mechanisms (Conway, 2020).

On the other hand, this work raises new questions for future research. For instance, are the mechanisms underlying crossmodal statistical learning functionally and neurophysiologically comparable to those of unimodal statistical learning? Additionally, is crossmodal statistical learning more effective than unimodal learning when attentional biases are controlled?

## Author contributions

D.D., R.D.B., and A.P.B. designed the experiments. D.D. and F.G. conducted the experiment. A.P.B. analyzed the data. F.G. and A.P.B. wrote the original draft. R.D.B. and A.P.B. edited the manuscript. D.D. and F.G have contributed equally to the development of this project.

## Acknowledgments

We thank Alicia Macià-Mendoza for help in data collection for Experiment 3. This work was supported by the AGAUR (BP 00213), and the Spanish Ministerio de Ciencia, Innovación y Universidades, which is part of Agencia Estatal de Investigación (AEI), and the European Regional Development Fund awarded to the projects <u>RTI2018-100977-J-I00</u> and <u>RYC2022-037652-I</u> to A.P.B. and PID2021-127146NB-I00 project to R.D.B) and the Maria de Maeztu to the Institute of Neuroscience. The work was also supported by the SGR (2021 SGR 00352) from AGAUR, Generalitat de Catalunya to R.D.B. We also thank the CERCA Programme / Generalitat de Catalunya for institutional support.

