## Supplemental Figure 1 for "Predictable Modality Transitions and Amodal Representations Enable Crossmodal Statistical Learning"

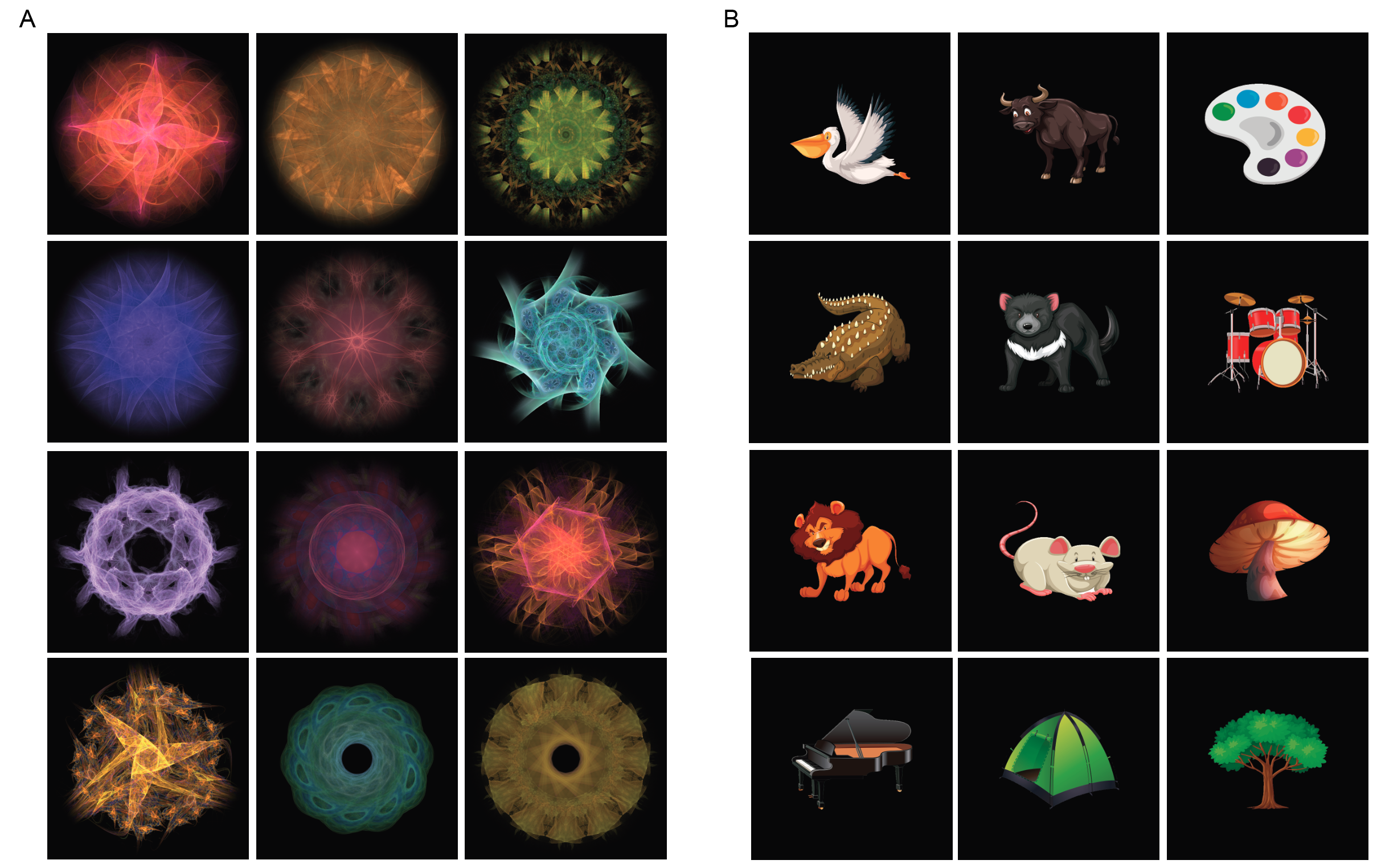

Fig S1 Synthetic fractals used as visual stimuli in experiment 1, 1b and 2. Drawings used in experiment 3 are copyrighted and cannot be published without permission. Here we show them partially masked.
